# Bacterial muropeptides promote OXPHOS and suppress mitochondrial stress in normal and human mitochondrial disease models

**DOI:** 10.1101/2023.01.05.522895

**Authors:** Dong Tian, Mingxue Cui, Min Han

## Abstract

Mitochondrial dysfunction critically contributes to many major human diseases but effective therapeutic methods to combat mitochondrial dysfunction are lacking^1,2^. Mitochondria of animal cells are sites of interaction between gut microbial metabolites and host factors, and many such interactions may be beneficial to mitochondrial health and host physiology^3^, albeit that specific beneficial interactions and the underlying mechanisms remain to be uncovered. Here we report the role of muropeptides derived from bacterial cell wall peptidoglycan (PG) in promoting mitochondrial functions in mammalian models. We found that muropeptides directly bind to ATP synthase, which stabilizes the complex and promotes its activity. The benefit is seen in increased oxidative respiration and mitochondrial membrane potential, as well as decreased oxidative stress in human intestinal epithelial cells (IECs). Strikingly, we also found that muropeptide treatment can recover mitochondrial structure and functions, as well as inhibit several pathological phenotypes of mutant fibroblast cells derived from mitochondrial disease patients. In mice, we show muropeptides accumulate in mitochondria of IECs and promote small intestinal homeostasis and nutrient absorption by modulating energy metabolism. This study identifies ATP synthase as a muropeptide receptor and the corresponding physiological function in mammals, and points to a potential treatment for human mitochondrial dysfunction diseases.

**Highlights:**

- Bacterial muropeptides bind and stabilize ATP synthase complex and enhance its activity
- Muropeptides enhance mitochondrial OXPHOS in human IECs
- Muropeptides recover mitochondrial structure and functions in mitochondrial disease cells
- Muropeptides promote small intestinal epithelial homeostasis and nutrient uptake in mice

## Introduction

Extensive studies in past decades have indicated that mitochondrial (Mt) dysfunction, commonly marked by dysfunction in oxidative phosphorylation (OXPHOS), critically contributes to the pathogenesis of many major human diseases known as Mt diseases ^1,2^. More specifically, OXPHOS dysfunction is known to result in various clinical manifestations, including many neurological disorders, cardiomyopathies, metabolic diseases, and others ^2,4-6^. For example, Leigh syndrome (LS), the most common childhood-onset disease among a large number of genetic Mt diseases, can be caused by mutations in 75 different genes, most of which encode proteins acting in OXPHOS ^7^. In addition, effective methods to suppress OXPHOS dysfunction are highly sought after as potential treatments for many Mt diseases including neurodegenerative diseases.

The Mt is known to be the site for interactions between microbial metabolites and host factors. While most previous studies focused on harmful interactions between pathogenic bacteria and host Mt^8^, some studies have suggested that Mt are an important hub for microbiota-host crosstalk that beneficially affect host physiology ^3,9^, which is consistent with the symbiosis hypothesis. However, specific beneficial roles on Mt functions, especially on OXPHOS function, by specific microbial metabolites and the underlying mechanisms remain largely unexplored.

Peptidoglycan (PG) is a crucial and unique component of the bacterial cell wall in both Gram-positive and Gram-negative species^10^. PG derivatives are among the microbial compounds released into the intestinal lumen of the host. Specifically, up to 50% of PG is degraded and released by hydrolysis that generates PG fragments of diverse structures known as muropeptides^11,12^. In mammals, cytosolic NOD-like receptors (NOD1 and NOD2) and peptidoglycan recognition proteins (PGRPs) are the well-known muropeptide receptors ^11,13,14^. Muropeptides have been shown to translocate from intestinal cells to many tissues including the blood stream and brain to promote the host innate immune system and neuronal activity through interactions with these known receptors ^13,15-18^. However, the role of muropeptides in host Mt was not known, and given the potentially high abundance of muropeptides in intestinal epithelial cells, muropeptides may benefit host physiology through unknown receptors in mammals. Such a possibility was suggested by our recent finding that muropeptides suppress Mt stress and promote animal development in the nematode *C. elegans* and do so by entering Mt and interacting with the ATP synthase ^19^.

Since bacteria are food for *C. elegans*, the conservation of a muropeptide-Mt interaction^19^ and the functional significance in mammals may not be automatically assumed despite the structural conservation of ATP synthase. Given the importance of finding a potential mechanism to combat OXPHOS dysfunction, we explored this novel muropeptide function in mammals. In this study, we show that muropeptides enter the Mt of mammalian IECs and promote Mt homeostasis. These muropeptides directly bind to multiple subunits of ATP synthase, stabilize the enzyme complex, and promote its activity. Consequently, cells treated with muropeptides have drastically higher oxidative respiration and membrane potential, as well as decreased ROS levels. These beneficial roles of muropeptides were also prominently observed in mammalian LS disease models with OXPHOS dysfunction. Moreover, we show that muropeptides regulate Mt energy metabolism and promote small intestinal epithelial homeostasis and nutrient absorption in mice. This work uncovers a microbiota-Mt communication modality involved in small intestinal homeostasis and function, and a potential therapeutic treatment for Mt diseases.

## Results

### Microbial muropeptides accumulate in mammalian mitochondria, stabilize the ATP synthase complex, and enhance its activity

During cell division, PG in bacteria is hydrolyzed by the combination of many bacterial enzymes into PG fragments that are also known as muropeptides^11,12^. Larger PG molecules released from intestinal bacteria may also be digested into muropeptides by host-secreted lysozymes ^19,20^. To investigate if muropeptides have potential beneficial roles in promoting Mt functions in the mammalian small intestine, we first tested if muropeptides are found in the Mt of mouse small intestinal epithelial cells (IECs). Applying the HEK-Blue-NOD1 assay system^21^ on Mt lysate from IECs isolated from mice, we found that muropeptides accumulated at a significant level in the Mt of mouse IECs (Fig. 1a). We then showed that *E. coli*-derived PG muropeptides also bind to ATP synthase from human IECs (caco-2 cells) in a concentration dependent manner (Fig. 1b), which is consistent with our finding in *C. elegans* ^19^. The short peptide feature of the muropeptide structure was required for binding, as protease K, but not trypsin, treatment of the PG muropeptides eliminated the binding (Fig. 1c). ATP synthase is a large complex comprised of at least 22 subunits ^22^. To better understand how muropeptides interact with ATP synthase, we purified recombinant subunits of the hydrophilic F1 subcomplex from both *C. elegans* and *H. sapiens* (Extended Data Fig. 1a, b) and performed PG muropeptide pull-down assays. The collective result indicated that PG muropeptides directly bind to F1 subunits α, d and F6, but not subunit β (Fig. 1d, e).

**Fig 1.**
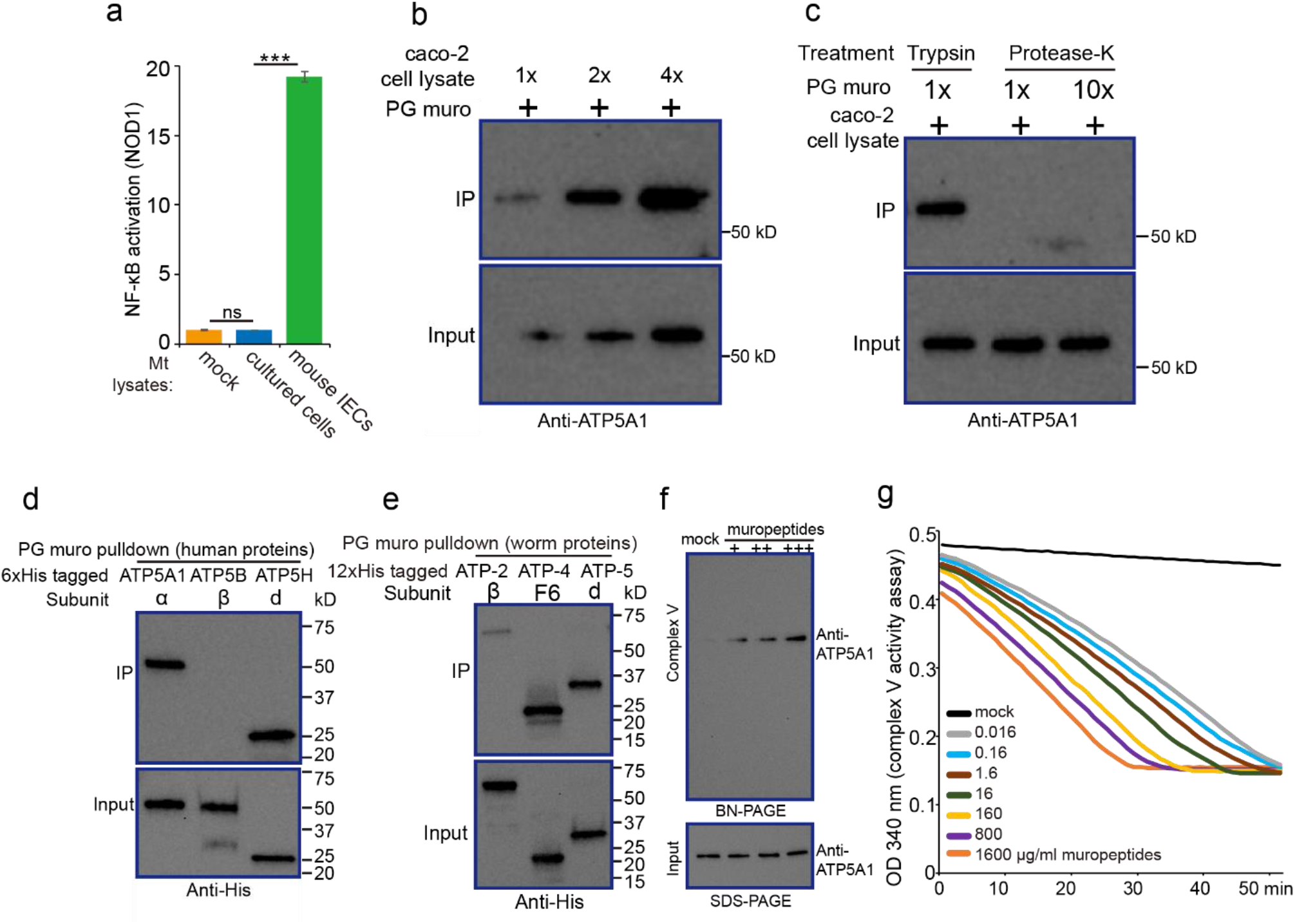
Muropeptides stabilize ATP synthase and promote its activity directly. **a**, Bar graph showing NF-κB activation that indicates the interaction between muropeptides from Mt lysates of intestine-derived mammalian cells and the NOD-1 receptor. Mt lysate from mouse IECs induced dramatically higher NF-κB activity than the lysate from MODE-K cultured cells. Mean ± SEM from 3-4 replicates. **b**, Interaction between PG muropeptides and human ATP synthase α subunit (ATP5A1) detected by *in vivo* pull-down assays. ATP5A1 from caco-2 cells bound to PG muropeptides, and the binding increased with more lysate input. **c**, Treatment with Protease-K, but not trypsin, eliminated the binding of PG muropeptides to ATP5A1. **d-e**, PG muropeptides specifically bound to the α, d and F6 subunits, but not β subunit, of mammalian (d) or worm (e) ATP synthase. **f**, BN-PAGE analysis showed that the ATP synthase Complex (Complex V) in Mt lysates from caco-2 cells was increased by adding muropeptides. Immunoblotting performed with anti-ATP5A1 antibody. SDS-PAGE electrophoresis and immunoblotting of ATP5A1 was used as the loading control. +: 5 ng/ml; ++: 500 ng/ml; +++: 500 μg/ml muropeptides. **g**, Complex V activity increased with increased muropeptide concentration. Data are averaged from 3 replicates. Inserted box: a simplified schematic representation of reaction that triggered by complex V. *** p<0.001 from Student’s t tests.

Based on these results, we hypothesized that muropeptides may promote ATP synthase activity by binding to the F1 subcomplex and stabilizing the enzyme complex (complex V). Indeed, analysis using non-denaturing blue native BN-PAGE revealed that the level of intact complex V increased with addition of muropeptides to soluble mitochondrial lysates from caco-2 or bovine heart cells (Fig. 1f; Extended Data Fig. 1c, d). Importantly, we found that muropeptides could directly increase ATP synthase activity in a concentration dependent manner by a standard complex V activity assay (Fig. 1g). Together, these findings indicate that muropeptides bind, stabilize, and enhance the activity of mammalian ATP synthase.

### Muropeptides promote OXPHOS in human cultured IECs

To assess whether muropeptides promote Mt function in mammalian IECs, we analyzed the Mt oxidative respiration of caco-2 cells, a cell line derived from human colon cancer that morphologically and functionally resembles enterocytes ^23^. By measuring the oxygen consumption rate (OCR)^24^ with a Seahorse XFe24 Analyzer, we observed that muropeptide treatment caused a significant increase in the basal, ATP-linked, and maximal mitochondrial OCR in caco-2 cells (Fig. 2a, b). We also found that muropeptide treatment boosted the ATP level and Mt membrane potential (Δψ_m_) in these cells (Fig. 2c, d). Furthermore, consistent with our finding in *C. elegans* ^19^, muropeptide treatment also inhibited ROS production in caco-2 cells (Fig. 2e), supporting a conserved function of muropeptides in promoting Mt oxidative respiration and suppressing oxidative stress. In addition, we showed that muropeptide treatment did not increase the protein level of ATP synthase α subunit (Extended Data Fig. 2), suggesting that the increase of Mt oxidative respiration was not due to an increase in the cellular number of ATP synthase. Altogether, these data indicate that muropeptides are ATP synthase agonists that significantly boost ATP production and OXPHOS in the Mt of cultured human IECs. This indication may also suggest that “under-performing” ATP synthase has limited the capacity of OXPHOS in these cells without muropeptides.

**Fig. 2.**
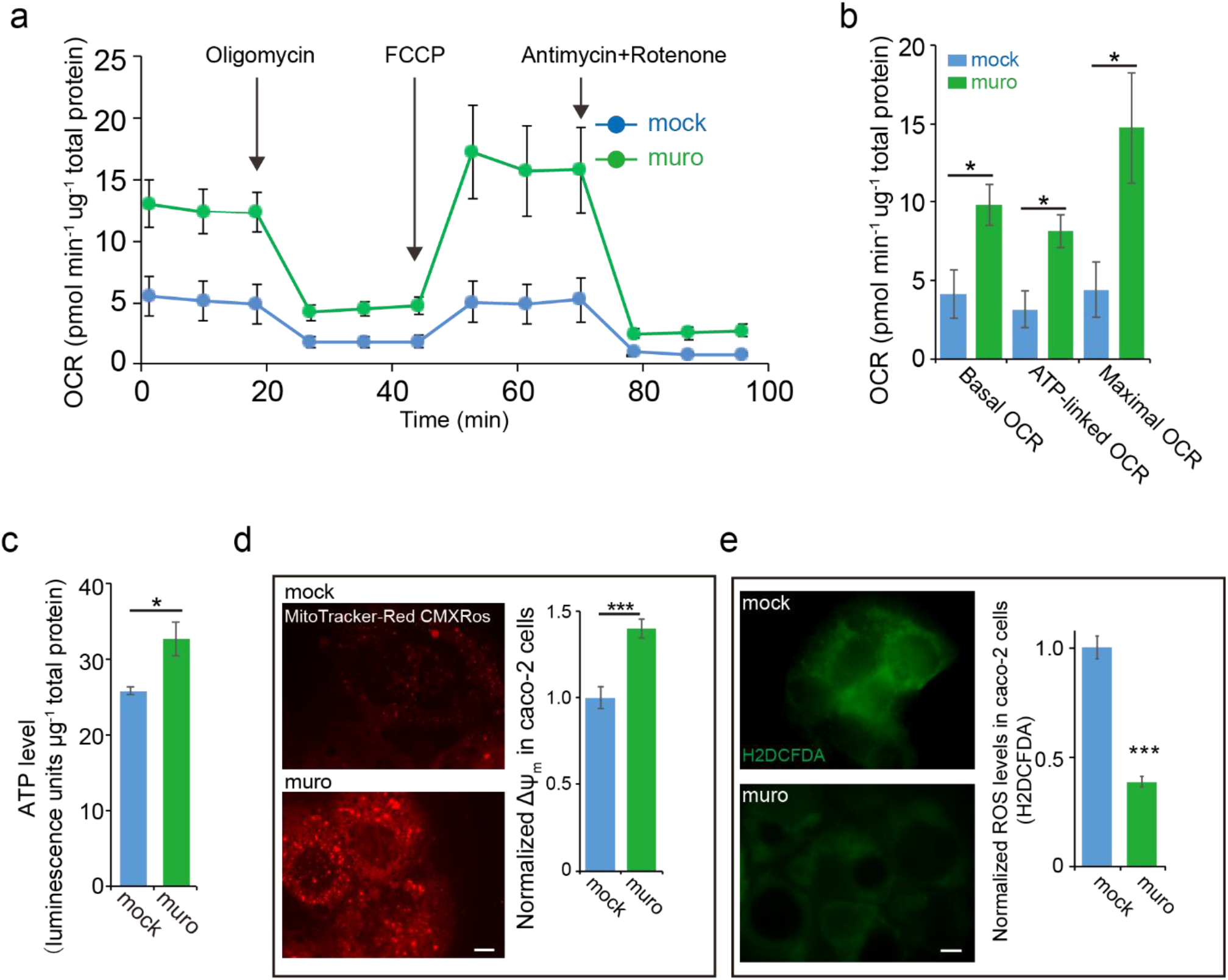
Muropeptides enhance mitochondrial function in human IECs. **a**, Data from a Mt oxidative respiration analysis of caco-2 cells show the increase in oxygen consumption rate (OCR) by supplementation with muropeptides (800 μg/ml). Oligomycin (2.5 μM), an ATP synthase inhibitor, was added to evaluate the ATP-linked increase in OCR. FCCP (1.0 μM), an uncoupling agent that raises OCR to maximal value by disrupting the Mt proton gradient and membrane potential, was added to measure maximal OCR. Antimycin and rotenone (1.25 μM), two inhibitors of the Mt respiration chain, were added to evaluate non-Mt OCR. The “mock” sample consisted of the same cells supplemented with lysozyme solution. Mean ± SEM from 3 replicates. **b**, Bar graphs of the OCR quantification show that the majority of the increase in OCR in caco-2 cells by muropeptide (800 μg/ml) supplementation was due to increase in ATP-linked or Mt-related OCR. Basal OCR: last OCR measurement before oligomycin injection. ATP-linked OCR: basal OCR minus minimal OCR value after oligomycin injection. Maximal OCR: maximum OCR measurement after FCCP injection. Mean ± SEM from 3 replicates. **c**, Bar graph shows significant increase of ATP level in caco-2 cells supplemented with muropeptides (800 μg/ml). Mean ± SEM from 3 replicates. **d**, Microscope images and bar graphs showing that addition of muropeptides (800 μg/ml) to caco-2 cells increased Mt membrane potential (Δψm) as indicated by MitoTracker-Red CMXRos (red fluorescence). n = 108-126 cells, mean ± SEM. Scale bar =20 μm. **e**, Microscope images and bar graphs showing that addition of muropeptides (800 μg/ml) to caco-2 cells suppressed ROS production as indicated by H2DCFDA (green fluorescence). n=49-57 cells, mean ± SEM. Scale bar =20 μm. * p<0.05; *** p<0.001 from Student’s t tests.

### Muropeptides recover mitochondrial structure and promote mitochondrial function in OXPHOS-defective human cells

Since we found that muropeptides had a beneficial impact on “wild type” cells, we next tested if muropeptides could also benefit cells with Mt OXPHOS defects. Among a large number of so-called Mt dysfunction diseases, Leigh Syndrome (LS) is the most common and well described inherited neurodegenerative disorder that is directly caused by damage to OXPHOS, as it can be caused by mutations in a large number of genes acting in the Mt OXPHOS complex^7^. The disease is associated with multiple neuronal and neuromuscular dysfunctions including loss of motor skill, seizure, and heat failure^7,25^. We thus obtained LS mutant fibroblast cells (GM13411, Coriell Institute) which were isolated from a patient diagnosed with Leigh Syndrome (LS) and hypertrophic cardiomyopathy ^26^. GM13411 contains two mutations in two OXPHOS complexes. One is a T8993G mutation in the Mt DNA that leads to a Leu to Arg change at aa156 in the ATP6 subunit of ATP synthase (Extended Data Fig. 3a)^26^. The other is a frameshift mutation (79delC) in pyruvate dehydrogenase (PDHA1), which is a key subunit of the pyruvate dehydrogenase complex (PDHc) that catalyzes the conversion of pyruvate to acetyl-coA to drive the activities of the TCA cycle and electron transport chain (ETC)^27^. Both mutations may be critical drivers of LS pathogenesis (see Discussion).

We first confirmed that the GM13411 fibroblasts have severe oxidative phosphorylation defects, compromised ATP production, and increased ROS (Extended Data Fig. 3b-e), which is consistent with previous analysis of this cell line ^28,29^. Since we showed that muropeptides stabilized the mammalian ATP synthase complex and promoted the activity (Fig. 1f, g; Extended Data Fig. 1c, d), we hypothesized that muropeptides may also be able to significantly improve Mt oxidative respiration in mutant cells with lesions in at least some OXPHOS complexes. To test this hypothesis, we supplied muropeptides to GM13411 cells and found that Mt oxidative respiration was significantly restored (Fig. 3a, b). Consistent with this result, we also observed that muropeptide treatment boosted ATP production and dramatically reduced ROS production in these cells (Fig. 3c, d).

**Fig. 3.**
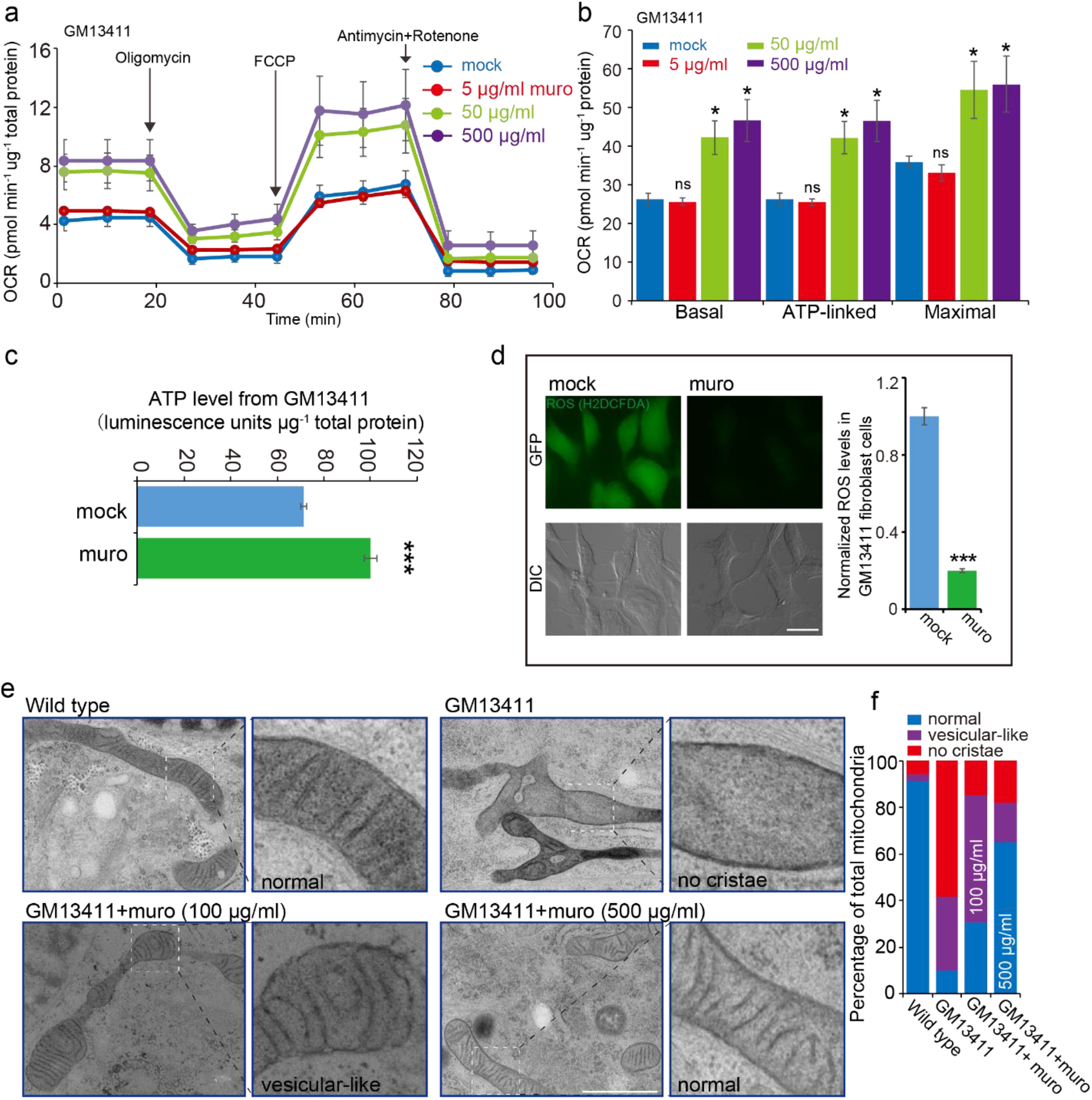
Muropeptides significantly recover mitochondrial function and structure of GM3411 mutant cells. **a**, Mt oxidative respiration analysis of GM13411 mutant fibroblasts showing that muropeptide supplementation increases oxygen consumption rate (OCR). The “mock” treatment is supplementation of lysozyme solution without PG. Mean ± SEM from 3 replicates. **b**, Bar graphs of the OCR quantification show that the basal OCR, ATP-linked OCR and maximal OCR in mutant fibroblast cells were increased by muropeptide supplementation. Mean ± SEM from 3 replicates. **c**, Bar graph showing significant increase of ATP level in GM13411 mutant fibroblasts supplemented with muropeptides (500 μg/ml). Mean ± SEM from 3 replicates. **d**, Representative images and bar graph showing that muropeptide (50 μg/ml) supplementation reduced ROS levels in GM13411 mutant fibroblasts as indicated by H2DCFDA dye staining. n=64-68, mean ± SEM. Scale bar in (d), 20 μm. **e** and **f**, Representative electron microscopy (EM) images (e) and statistical analysis (f) of Mt structures in WT, GM13411 mutant, and muropeptide-treated mutant fibroblast cells. While most of Mt in GM13411 fibroblast cells show no cristae (top right panel), muropeptide supplementation recovers cristae structure in mutant fibroblasts in a concentration-dependent manner (bottom panels). “Vesicular-like” indicates that Mt have cristae structure, but the cristae morphology is still abnormal. n=68-214 Mt from >10 micrographs. Scale bar in (e) is 100 nm. *** p<0.001, * p<0.05, ns: not significant.

Severe damage in OXPHOS complexes is known to disrupt Mt structure^30,31^. We then analyzed the Mt structure of GM13411 cells with or without muropeptide treatment by electron microscopy. While most of the Mt from GM13411 cells lacked cristae structure that are commonly found in wild-type Mt, muropeptide treatment significantly recovered the cristae structure in the mutant Mt in a concentration dependent manner (Fig. 3e, f).

Overall, these results indicate that muropeptide treatment can effectively recover OXPHOS activity and Mt structure, as well as suppress oxidative stress, in LS fibroblasts with severe defects in OXPHOS.

### Muropeptides suppress mitochondrial stress and promote fitness of LS cells

Mt stress and dysfunction are known to disrupt the structure and function of lysosomes and thus cause an increase in the accumulation of vacuoles and autophagosomes ^32-34^. Indeed, aberrant accumulation of autophagosomes and vacuoles were observed in GM13411 cells, and such an increase was partially suppressed by muropeptide supplementation (Fig. 4a, b; Extended Data Fig. 4a, b). In addition, muropeptide treatment drastically reduced the expression of Mt stress marker hsp60 (Fig. 4c, d) as well as autophagy marker LC3-II (Fig. 4e, f).

**Fig. 4.**
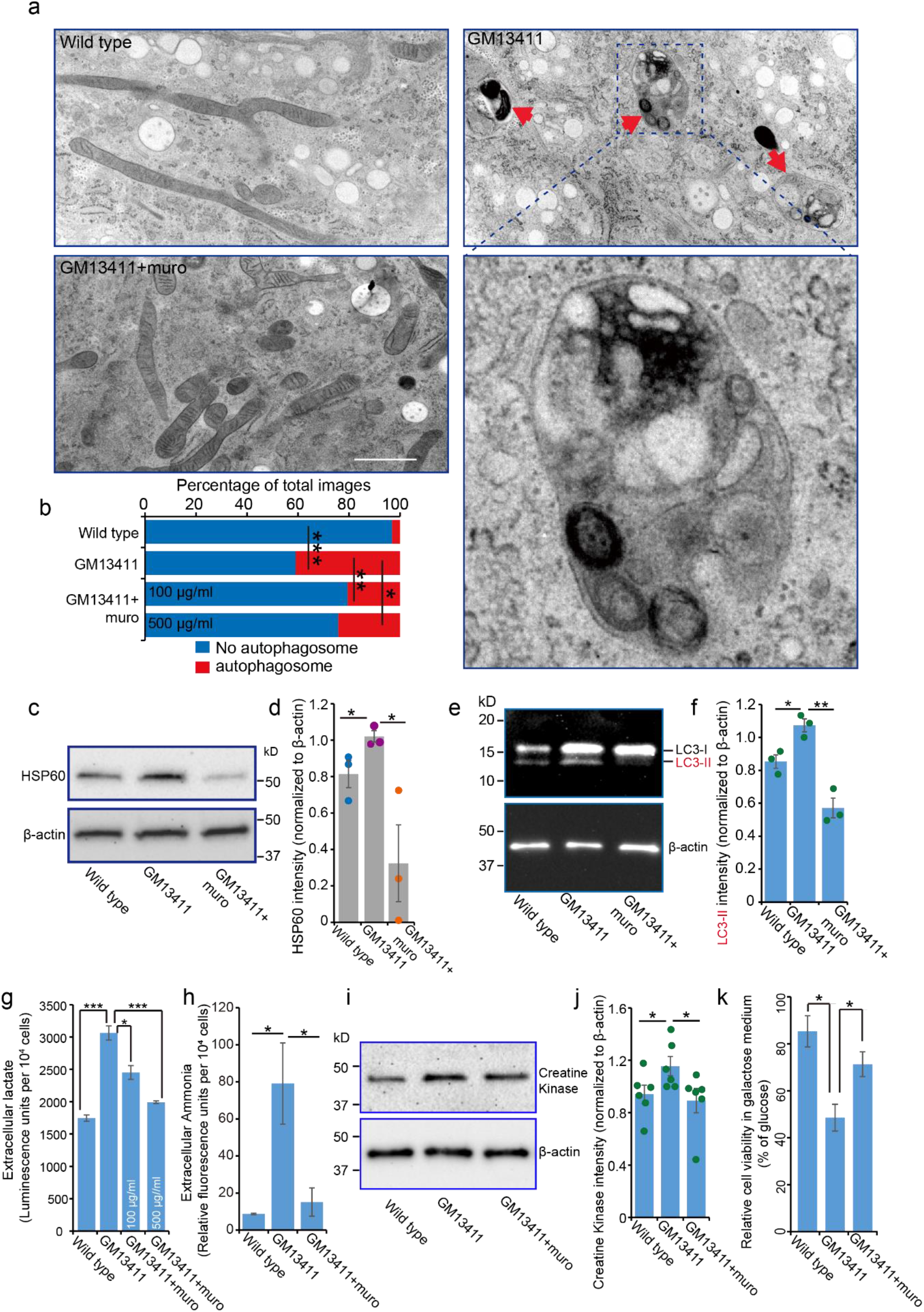
Muropeptides promote the fitness of GM13411 mutant cells. **a** and **b**, Representative EM images (a) and quantification (b) show that autophagosomes (red arrows) are enriched in GM13411 mutant cells and muropeptide supplementation decreased this accumulation. Scale bar in (A); 1 μm. **c** and **d**, GM13411 mutant fibroblasts have increased HSP60 expression, a marker of mitochondrial stress, and muropeptide supplementation dramatically decreased HSP60 protein level. Mean ± SEM from 3 replicates. **e** and **f**, GM13411 mutant fibroblasts have increased autophagy marker protein LC-3II expression, which was dramatically decreased by muropeptide supplementation. Mean ± SEM from 3 replicates. **g**, GM13411 mutant cells produced more lactate, which is the end-product of glycolysis, compared to wild type cells. Muropeptide supplementation significantly decreased lactate levels. Lactate levels were measured from media after 24 h culture and muropeptide treatment. Mean ± SEM from 3 replicates. **h**, GM13411 mutant cells generated more ammonia, which was suppressed by muropeptide (50 μg/ml) supplementation. Mean ± SEM from 3 replicates. **i** and **j**, GM13411 mutant fibroblasts have increased creatine kinase expression, which was decreased by muropeptide (500 μg/ml) supplementation. Mean ± SEM from 6 replicates. **k**, Cell viability in galactose medium was measured. GM13411 mutant cells exhibited low viability in galactose medium, which was partially rescued by muropeptide (50 μg/ml) supplementation. Data was normalized to that under normal glucose condition. Mean ± SEM from 3 replicates. *** p<0.001, ** p<0.01, * p<0.05.

Many LS patients, including the one from which the GM13411 cells were derived, were diagnosed with elevated levels of creatine kinase, lactate, and ammonia in the blood, indicating changes in energy metabolism and reduced fitness of the cells that are the consequence of decreased OXPHOS and increased glycolysis ^26,35^. The GM13411 fibroblasts recapitulated these pathological features (Fig. 4g-j; Extended Data Fig. 4c, d). Our analysis showed that muropeptide treatment significantly reduced the levels of all three molecules in GM13411 mutant cells (Fig. 4g, j; Extended Data Fig. 4c, d), indicating that muropeptides also improved the metabolism and fitness of mutant cells. In line with this idea, we also found that the cell viability was partially rescued by muropeptide treatment (Fig. 4k).

Leigh syndrome can be caused by mutations in many genes with different Mt functions^7^. We also tested a second LS patient-derived fibroblast cell line that does not have the lesion in ATP synthase (GM03672, Coriell Institute). GM0372 has also been shown to contain two mutations in two different OXPHOS complexes. One is the exact same frameshift mutation in PDHA1 of the PDHc as that in GM13411 and the other is a mtDNA point mutation in MTCYB (cytochrome b, subunit of Complex III) (Extended Data Fig. 5a) ^27,36^. The deficiency of PDHc in these cells was previously determined ^37^. Similar to GM13411, GM0372 also exhibited elevated levels of ROS and lactate, and reduced survival rate (Extended Data Fig. 5b-e). We then found that muropeptide treatment significantly suppressed all of these defects (Extended Data Fig. 5b-e). The results above support the notion that, by promoting OXPHOS and suppress Mt stress, muropeptides significantly recover the fitness of LS mutant cells.

### Muropeptides have a profound impact on ROS level and Mt energy metabolism in mouse IECs

Since we identified a high level of muropeptides in Mt extracted from mouse IECs (Fig. 1a), we next investigated the role of muropeptides in the small intestine of mice. To do this, we first used an antibiotic cocktail to deplete the intestinal microbiota in order to eliminate the source of muropeptides, following a well-established protocol for antibiotic induced microbiome depletion (AIMD) ^38^(Extended Data Fig. 6a-d). The procedure produced germ free-like phenotypes, such as enlarged cecum and reduced spleen size (Extended Data Fig. 6e-j)^39,40^, in both male and female mice.

AIMD treatment caused an increase in ROS levels in IECs isolated from both male and female mice, with a more dramatic increase observed in males (>10 fold) (Fig. 5a, b). Strikingly, the ROS increase in both male and female mice was mostly suppressed by PG muropeptide treatment through oral gavage (Fig. 5a, b); the PG molecules were presumably further hydrolyzed to muropeptides by lysozymes in the host intestine ^19^. This result indicates that muropeptides can inhibit Mt oxidative stress in mouse IECs *in vivo*. Consistent with the antagonistic impacts of AIMD and muropeptides on OXPHOS, we found that AIMD treatment caused a decrease in ATP level in male mice and the decrease was significantly suppressed by feeding PG (Fig. 5c). Intriguingly, the difference in ATP level between the three groups of IEC samples from female mice were too small to reach any statistical significance (Fig. 5d), suggesting that neither AIMD nor PG muropeptides significantly affect ATP production in female mice. Since the effects of both AIMD and muropeptides were stronger in male mice, some of our further tests focused mainly on male mice.

**Fig. 5.**
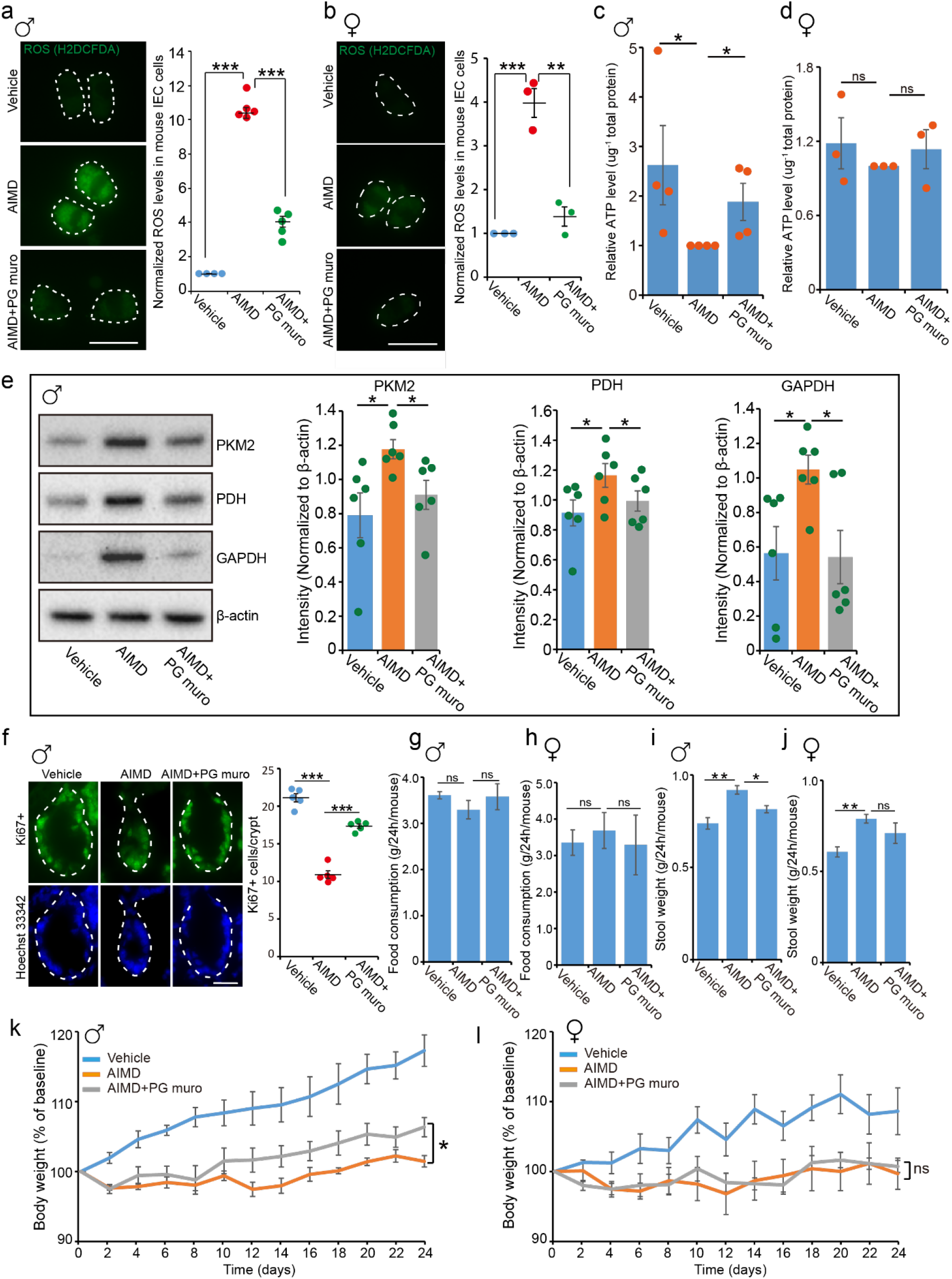
PG muropeptides promote small intestinal epithelial homeostasis and function. **a** and **b**, Representative images and quantification show ROS levels in small intestinal epithelial cells (IECs) of vehicle, AIMD or AIMD+PG muropeptide treated male (a) or female (b) mice. Mean ± SEM from 4-5 mice. Scale bar =20 μm. **c** and **d**, Bar graphs show ATP level in in small IECs of vehicle, AIMD or AIMD+PG muropeptide treated male (c) or female (d) mice. Mean ± SEM from 4 mice. **e**, Representative images and quantification showing that the expression of glycolytic proteins were increased in IECs from AIMD-treated male mice and the increase was suppressed with PG oral gavage. Mean ± SEM from 6 mice. **f**, Representative images and quantification showing proliferating cells (Ki67+) in small intestinal crypts were decreased in AIMD-treated male mice and the decrease was suppressed by PG oral gavage. Mean ± SEM from 5 mice. Scale bar =20 μm. **g** and **h**, Quantification showing that AIMD and/or PG muropeptide oral gavage has no significant effect on food consumption. Mean ± SEM from 4 mice per 24h. **i** and **j**, Quantification showing stool weight was increased in AIMD-treated male mice and the increase was partially suppressed by PG oral gavage. Mean ± SEM from 4 mice per 24h. **k** and **l**, Male (k) and female (l) mice weight gain during AIMD and/or PG muropeptide oral gavage. *** p<0.001, ** p<0.01, * p<0.05. ns: not significant. AIMD: Antibiotics induced microbiota depletion.

We then further analyzed energy metabolism that is known to be impaired by Mt oxidative stress ^41^. Indeed, while we found that expression of proteins in the glycolytic pathway were increased in IECs of AIMD-treated mice, they were strongly suppressed by PG oral gavage (Fig. 5e). These data suggest that muropeptides act to promote aerobic energy metabolism in the host small intestine.

### Muropeptide-Mitochondria interaction regulates small intestinal homeostasis and function in mice

Previous studies have linked Mt function to intestinal epithelial cell homeostasis ^42^. To determine the potential physiological consequences of muropeptides in the mouse small intestine, we examined proliferating cells in the small intestinal crypts. Consistent with previous findings ^39,43^, AIMD treatment drastically reduced proliferating cell numbers (Fig. 5f). Interestingly, we found that PG oral gavage strongly suppressed this reduction induced by AIMD treatment (Fig. 5f), indicating that muropeptides provide an important benefit to the homeostasis of the small intestine.

The small intestine is well-known as the primary site for nutrient sensing and absorption ^44^. To further assess whether muropeptides modulate nutrient absorption, we monitored food consumption and stool output in AIMD-treated mice. Although neither AIMD nor PG feeding caused statistically significant changes in food consumption in 24 hours (Fig. 5g, h), AIMD-treated mice produced more stool during this period (Fig. 5i, j), suggesting that microbiota depletion compromised digestion and nutrient uptake. Importantly, we found oral gavage of PG partially but significantly suppressed this increase in stool output in male mice, but not in female mice (Fig. 5i, j). While the difference between the two sexes is consistent with our observed difference in ATP level (Fig. 5c, d), the suppression effect in males suggests that PG muropeptides promote nutrient absorption and may do so by regulating Mt metabolism and small intestinal homeostasis. In line with these results, we found that AIMD treatment suppressed weight gain in both male and female mice, and weight gain was partially rescued by PG oral gavage in only male mice (Fig. 5k, l). To confirm that the muropeptide-Mt interaction is involved in this regulation, we tested Mt inhibitor oligomycin in male mice and confirmed that oligomycin oral gavage compromised Mt function in small intestine (Extended Data Fig. 7a-c). Importantly, oligomycin oral gavage also disrupted the homeostasis of small intestine, increased cecal weight and increased stool output, but not food consumption (Extended Data Fig. 7d-g). Consistently, oligomycin also reduced body weight in males (Extended Data Fig. 7h). All these data support that muropeptides mediate Mt metabolism responses for homeostasis and nutrient absorption in the mouse small intestine.

## Discussion

This study reveals a microbiota-Mt interaction mechanism by which peptidoglycan muropeptides from commensal bacteria enter Mt of host intestinal cells to promote energy metabolism and thus positively impact the homeostasis and physiological function of the host small intestine. Previous studies revealed diverse functions of PG muropeptides in the host, such as activating the immune system, regulating behaviors, and controlling body temperature of the host through the well-known receptors NOD1/2 or PGRPs ^13,15-18^. Here, we identified a new muropeptide receptor, the ATP synthase complex, in mammalian systems. Our mechanistic studies revealed that muropeptides could stabilize the ATP synthase complex and significantly raise its activity (Fig. 1f, g; Extended Data Fig. 1c, d). We further showed that this muropeptide-ATP synthase interaction enhanced Mt OXPHOS and energy metabolism, as well as other physiological functions in both cultured cells (Fig. 2-4; Extended Data Fig. 4, 5) and mice (Fig. 5; Extended Data Fig. 6, 7).

Since ATP synthase is a highly conserved enzyme and important therapeutic drug target ^45^, chemical molecules that could enhance ATP synthase activity would be highly desirable. However, although numerous inhibitors were discovered in the past decades, no chemical agonist of ATP synthase has been previously described in mammalian systems. The identification of muropeptides as ATP synthase agonists would provide not only a powerful tool to address mechanistic questions regarding Mt metabolism and homeostasis, but also a potential therapeutic drug for Mt dysfunction-related human diseases, including major neurodegenerative diseases. Such an attractive prospect is supported by the results from our tests in two LS disease cell lines, where muropeptides recovered OXPHOS activity, Mt structure, and other functions, as well as inhibited Mt stress and disease-related defects (Fig. 3, 4; Extended Data Fig. 3-5).

Results from our analyses under multiple cell conditions, including human IECs, fibroblasts from LS disease patients, and intestinal cells in mice, have indicated that increasing ATP synthase (Complex V) activity significantly boosts the overall OXPHOS activity including increases in membrane potential and reduction of ROS produced by electron transport chain (ETC) (Fig. 2a-e; 3a-d; 5a-c). These observations suggest that, without muropeptides, the relatively low ATP synthase activity has limited overall OXPHOS activity, including the activity of the ETC. By boosting ATP synthase activity, which includes pumping more protons back into the Mt matrix, muropeptides may also promote the activity of the ETC and thus increase the membrane potential. This action may be the mechanistic basis for their suppression of Mt dysfunction that results from defects in ETC.

The significant recovery of cristae structure (Fig. 3e, f) in GM13411 cells (LS disease fibroblasts) might suggest that such structural recovery was largely responsible for the increase in OXPHOS activity. Such a notion is consistent with previous studies suggesting that the assembly of intact ATP synthase on the inner membrane is significantly involved in generating cristae architecture^30,46^ and the GM13411 cells contain a Mt DNA missense mutation within the ATP6 subunit of ATP synthase, which was shown to disrupt the stability and activity of ATP synthase ^47,48^. However, several lines of evidence may suggest that muropeptides can promote OXPHOS through a mechanism that is independent of the increase in cristae structure or overall number of OXPHOS complexes. First, prominent increase in oxidative respiration and membrane potential, as well as drastic decrease in ROS, was also observed in cultured human IECs (caco-2) cells (Fig. 2) that are not expected to have the dramatic damage in cristae architecture associated with GM13411 cells. These changes are more likely due to increased proton translocation into the Mt matrix that promotes the earlier steps of OXPHOS. Second, a recent analysis of genomic DNA determined that GM13411 also contains a frameshift mutation in pyruvate dehydrogenase (PDHA1) which is a key subunit of the pyruvate dehydrogenase complex (PDHc)^27^. Since PDHc catalyzes the conversion of pyruvate to acetyl-coA that enters the TCA cycle to drive the activity in complex I and II, the PDHA1 allele is expected to significantly reduce oxidative respiration. Moreover, the second LS disease cell line, GM03672, has the exact same frameshift mutation in PDHA1 without the ATP6 gene mutation and has been shown to be deficient in PDHc activity ^27,37^. It is thus conceivable that PDHA1 is likely a key driver of the disease pathogenesis even though the ATP6 gene mutation may also be significantly involved in the case for GM13411. Such a notion is consistent with the findings that genetic mutations in 10 PDHc subunits are associated with LS^7^. Therefore, the rescue data from the tests in both LS cell lines (Fig. 3, 4; Extended Data Fig. 4, 5) are consistent with the idea that promoting ATP synthase activity may be able to suppress defects in other OXPHOS complexes. Finally, ETC dysfunction, which is accompanied with increased oxidative stress, is known to cause multiple defects in Mt structure, biogenesis and other functions^49,50^. Therefore, much of the structural defects in GM1341 cells, along with increased ROS, vacuoles, autophagy, lactate, ammonia, and creatine kinase, could be the consequence of disrupted ETC activities. The observed rescue by muropeptides may be mainly due to restored ETC activity.

In this study, we also analyzed the physiological functions of muropeptides in the small intestine of mice. Gut microbiota play important roles in intestinal homeostasis and function by regulating energy metabolism ^51-53^. Although several microbial metabolites have been discovered to promote energy metabolism and homeostasis, most studies focused on the large intestine ^44,54,55^. Here, we identified new bacterial metabolites, muropeptides, that promote small intestinal epithelial metabolism and proliferation through this muropeptide-ATP synthase interaction (Fig. 5a-f; Fig. 1b-g). Unlike the large intestine, the small intestine serves as the primary site of nutrient digestion and absorption. Some evidence suggests that microbiota are involved in nutrient absorption, exemplified by the observation that antibiotic-treated rats displayed reduced dietary lipid absorption ^56^ and conventionalized germ-free zebrafish had increased uptake of fatty acids under fasted and fed conditions ^57^. However, specific microbial metabolites and underlying mechanisms are still not clear. Using AIMD-treated mice and PG muropeptide oral gavage, we identified that bacteria-derived muropeptides could increase nutrient uptake in mice (Fig. 5g-j), suggesting a potential regulation of the host nutrient absorption by gut microbe-generated metabolites. In addition, our results indicated that muropeptides not only promote small intestinal homeostasis but also benefit other intestinal sections and even other organs such as the cecum and spleen (Extended Data Fig. 6e, g and j), suggesting a broad beneficial effect of muropeptides on host physiology and development.

## Acknowledgements

We thank Dr. Garry Morgan and Eileen O’Toole in the Boulder EM Services Core Facility at the Univ of CO for providing specimen preparation and imaging, Dr. Christopher C. Ebmeier in the Mass Spectrometry Facility at the Univ of CO for facilitating the proteomic analyses, Aileen Sewell for critical editing of the manuscript, and our lab members for providing helpful advice and discussion. This study was supported by NIH grants 1R35GM139631 (M.H.).

## Author contributions

D.T. designed research, performed experiments, analyzed the data, and wrote the paper; M.C. performed and analyzed data of some experiments; M.H. supervised the study and edited the paper.

---

**Extended Data Fig. 1.**
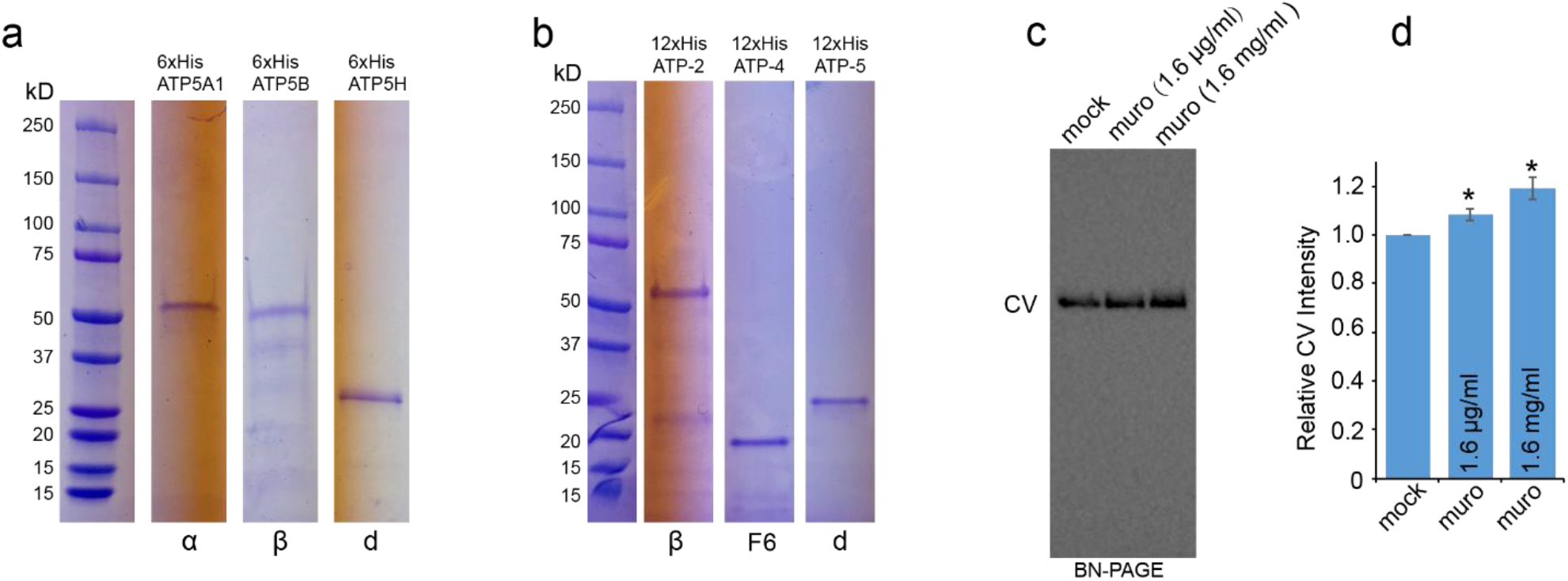
Muropeptides stabilize mammalian ATP synthase. **a** and **b**, Coomassie blue stained gel images show the purified recombinant ATP synthase subunits of human (a) and *C. elegans* (b). **c** and **d**, BN-Native PAGE analysis showing that the ATP synthase complex in Mt lysates from bovine heart were increased by adding increasing amounts of muropeptides. Immunoblotting performed with anti-ATP5A1 antibody. Mean ± SEM from 3 replicates. * p<0.05.

**Extended Data Fig. 2.**
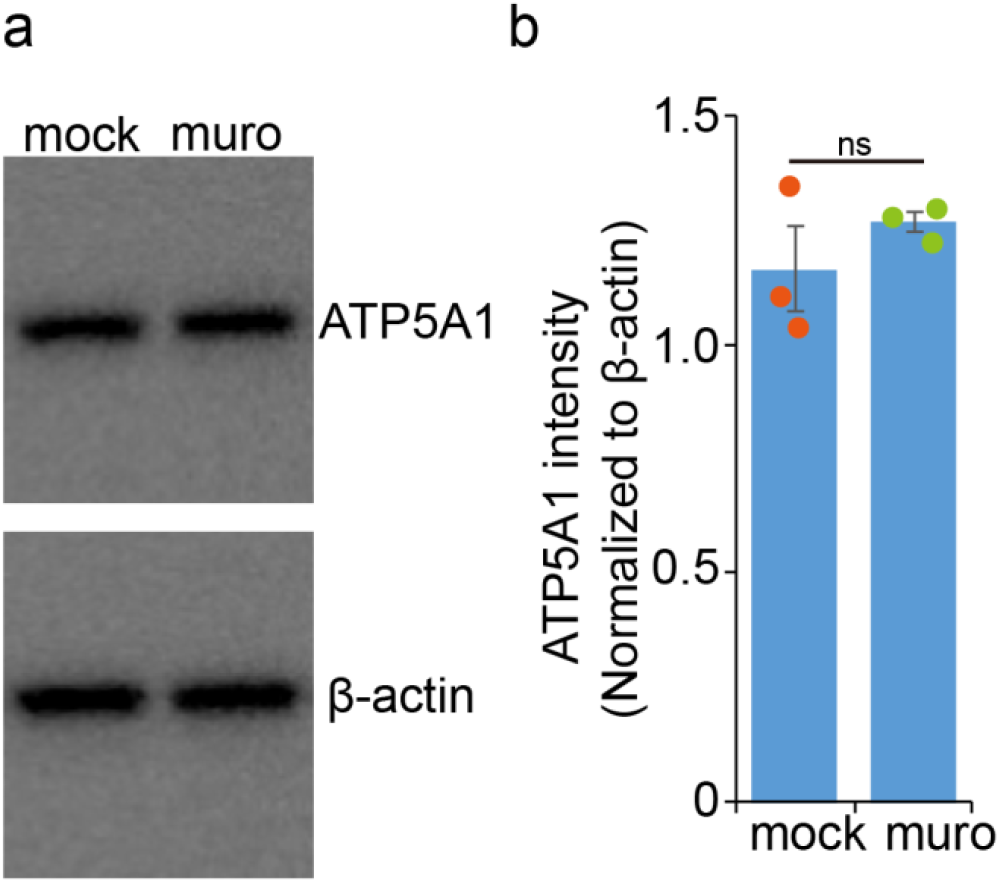
Muropeptides do not significantly change the protein level of the α subunit of ATP synthase. **a** and **b**, Western blot analysis of the protein level of the α subunit of ATP synthase in caco-2 cells supplemented with mock solution or muropeptides (800 μg/ml). The data were normalized to β-actin. Mean ± SEM from 3 replicates. ns: not significant

**Extended Data Fig. 3.**
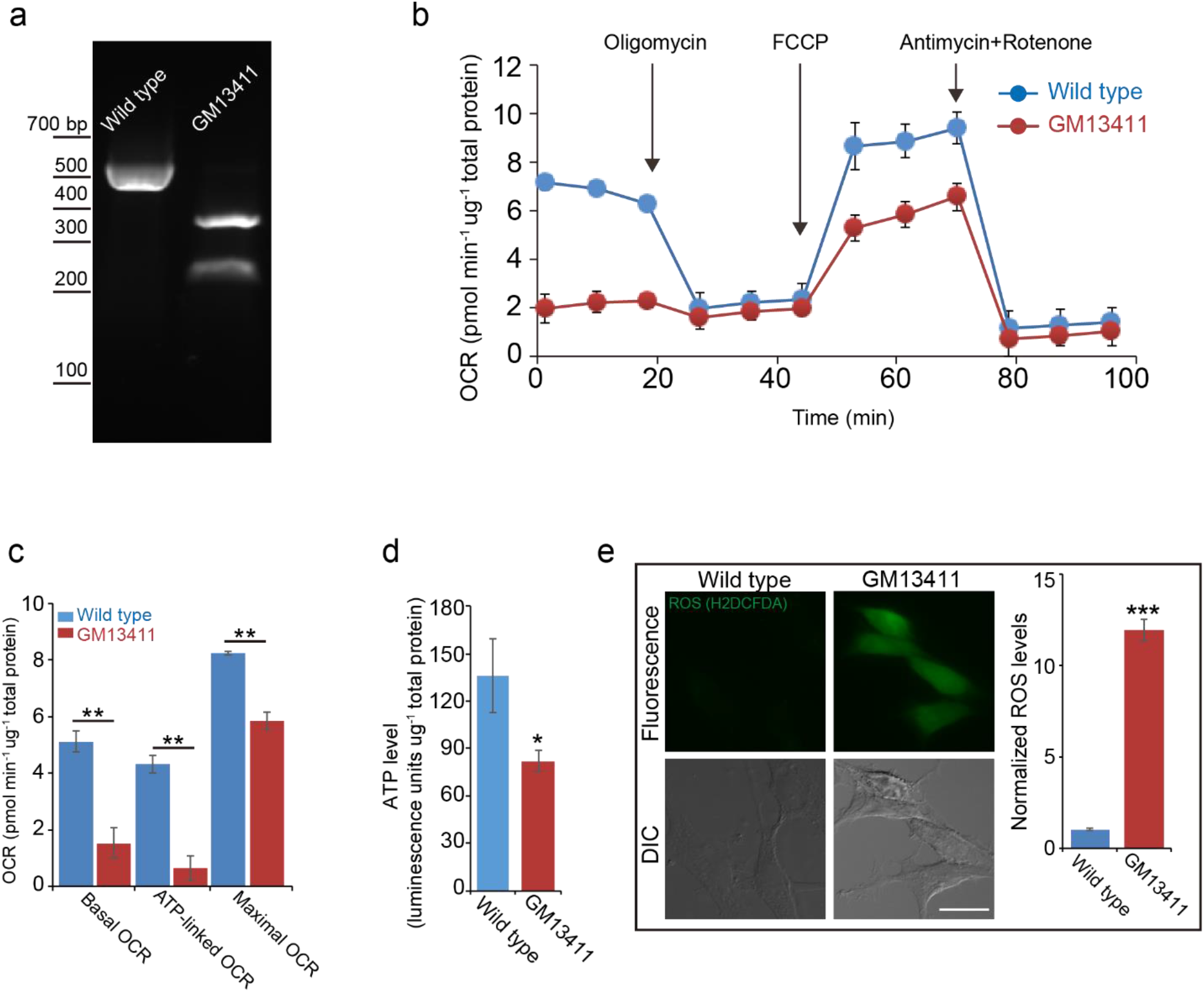
GM13411 mutant cells exhibit severe mitochondrial dysfunction. **a**, PCR genotyping of wild type and GM13411 mutant fibroblast cells. MspI was used to digest PCR fragments. **b**, Data from a Mt oxidative respiration analysis of wild type and GM13411 mutant fibroblast cells shows that the oxygen consumption rate (OCR) was decreased in the GM13411 mutant cells. Mean ± SEM from 3 replicates. **c**, Bar graphs of the OCR quantification show that basal OCR, ATP-linked OCR and maximal OCR were decreased in GM13411 mutant cells. Mean ± SEM from 3 replicates. **d**, Bar graph showing significant decrease of ATP level in GM13411 mutant cells, compared to wild type cells. Mean ± SEM from 3 replicates. **e**, Microscope images and bar graphs showing the increase of ROS as indicated by H2DCFDA (green fluorescence) in GM13411 mutant cells, compared to wild type cells. n= 53-56, mean ± SEM. Scale bar = 20 μm. *** p<0.001, ** p<0.01, * p<0.05.

**Extended Data Fig. 4.**
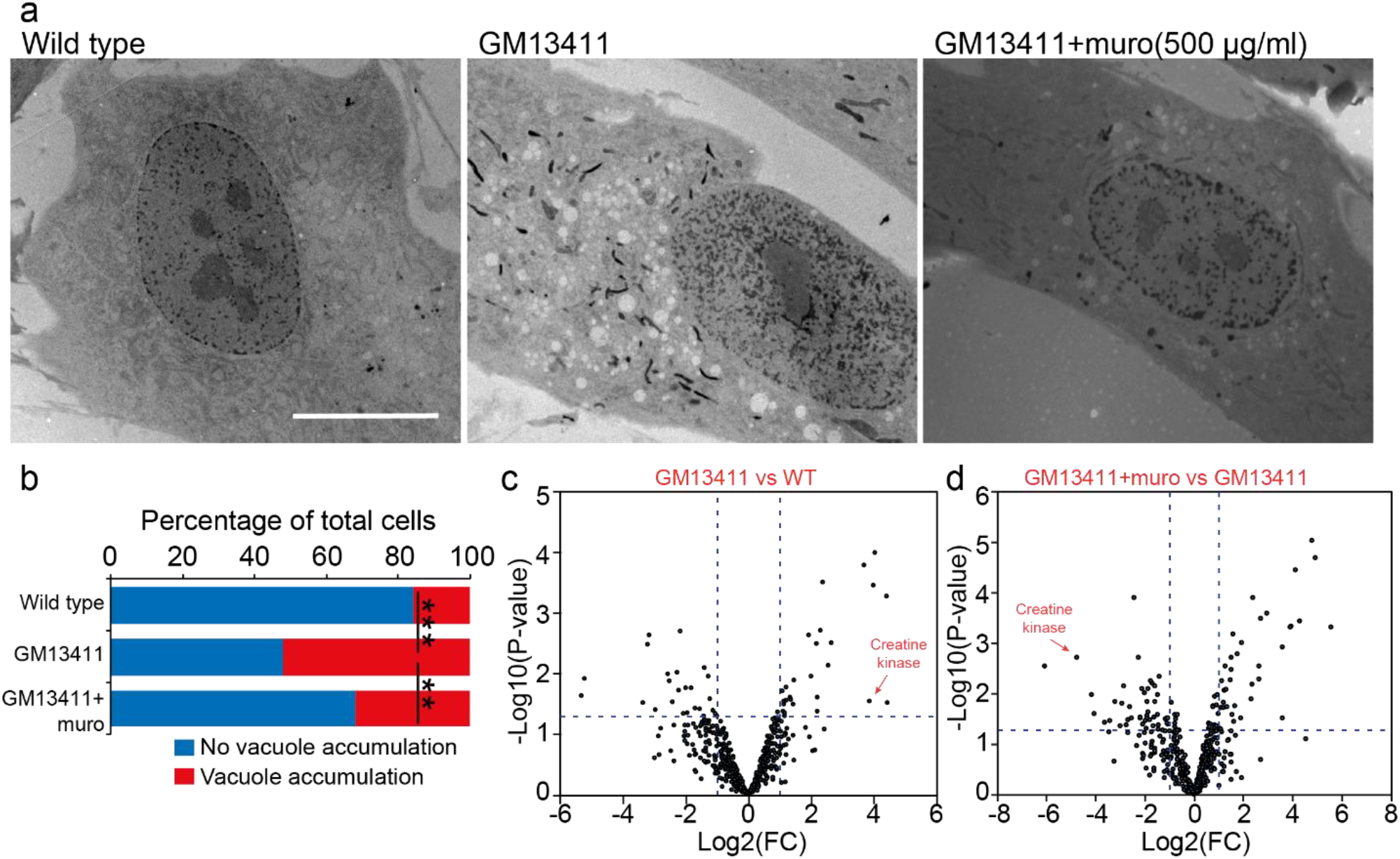
Muropeptides suppress vacuole accumulation and creatine kinase expression. **a** and **b**, Representative electron microscopy images (a) and quantification (b) showing that vacuole accumulation in GM13411 mutant cells was suppressed by muropeptide supplementation. Scale bar in (a) 10 μm. **c** and **d**, Volcano plots from puromycin immunoprecipitates and LC-MS analysis showing that creatine kinase protein synthesis was increased in GM13411 mutant fibroblast cells, compared to that in wild-type cells (c). Muropeptide supplementation (500 μg/ml) to GM13411 mutant cells significantly suppressed creatine kinase protein synthesis (d). *** p<0.001, ** p<0.0

**Extended Data Fig. 5.**
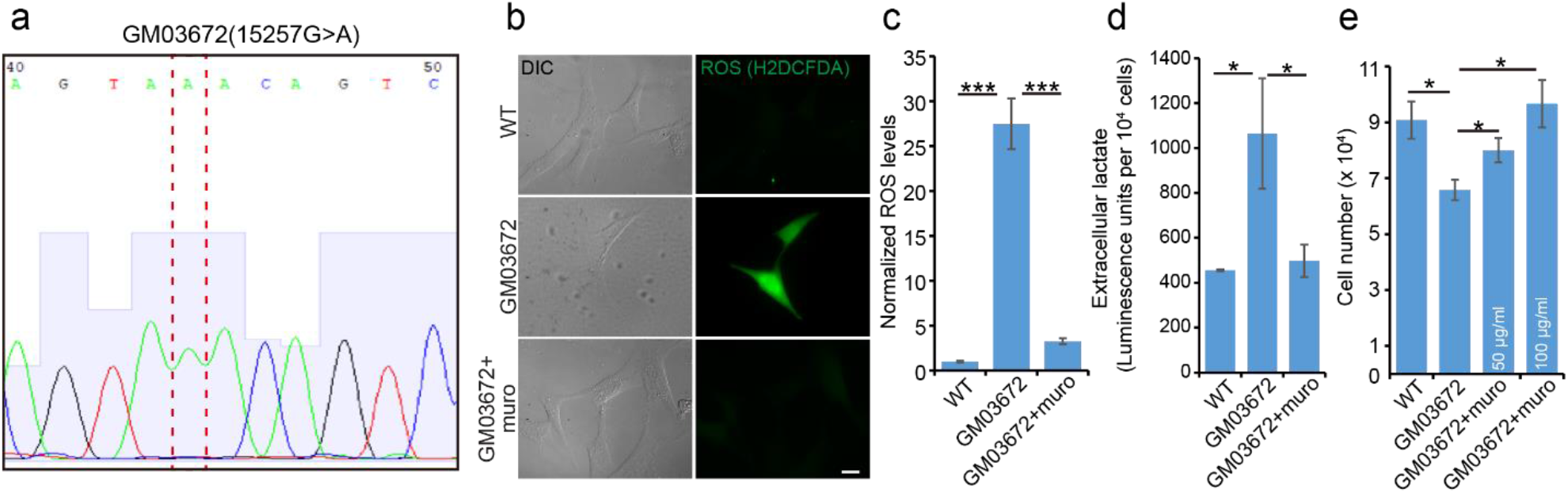
Muropeptides also suppress Mt dysfunction in another Leigh syndrome disease cell line. **a**, DNA sequencing of PCR fragment from GM03672 showing the 15257G>A point mutation. **b** and **c**, Representative images and bar graph showing that GM03672 mutant cells produced more ROS, which was suppressed by muropeptide (50 μg/ml) supplementation. n=64-78, mean ± SEM. Scale bar in (b), 20 μm. **d**, GM03672 mutant cells produced more lactate, compared to wild type control cells. Muropeptide supplementation (50 μg/ml) significantly decreased lactate levels. Mean ± SEM from 3 replicates. **e**, GM03672 mutant cells exhibited low viability, which was partially rescued by muropeptide supplementation. Mean ± SEM from 3 replicates. *** p<0.001, * p<0.05.

**Extended Data Fig. 6.**
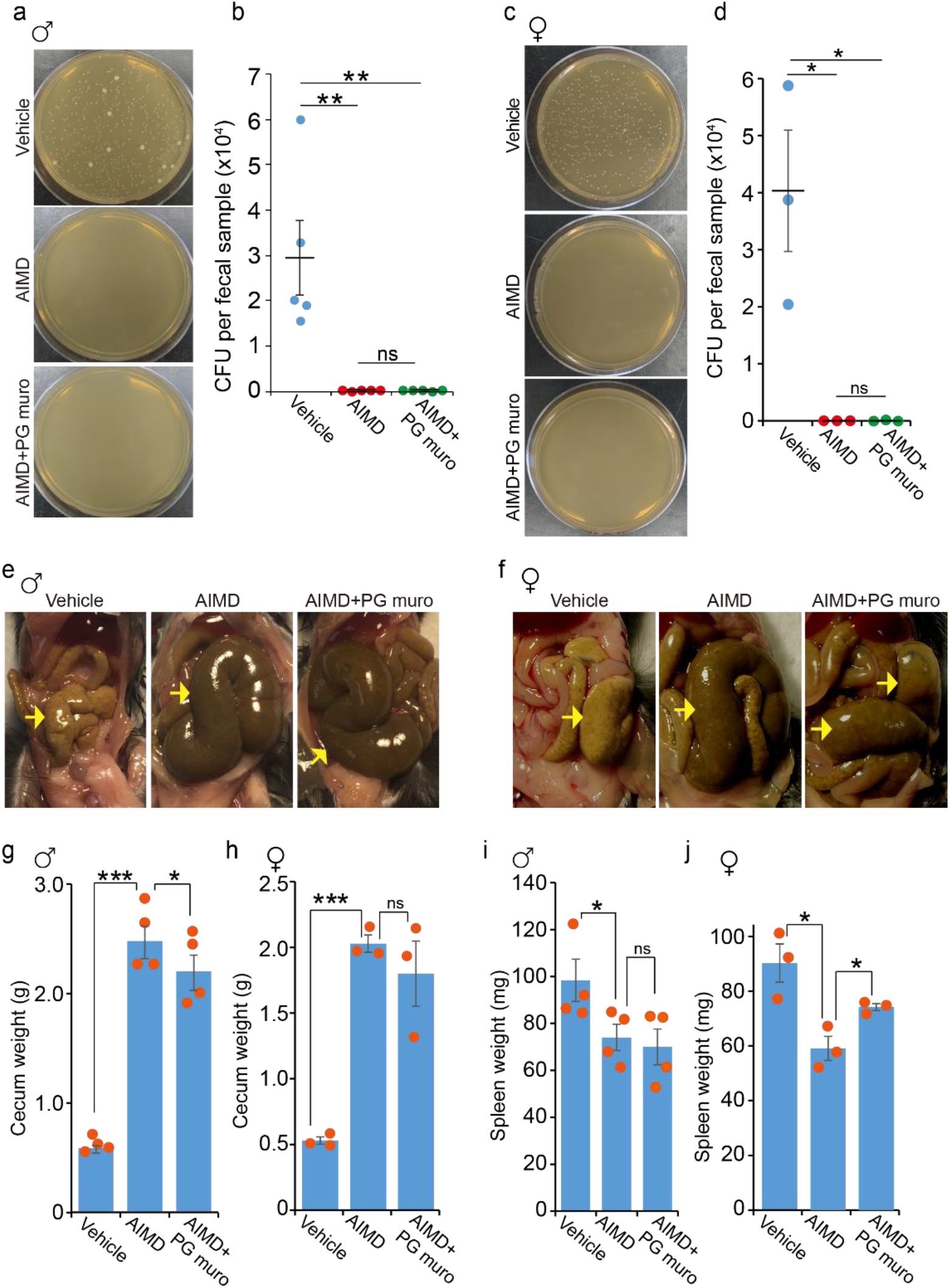
AIMD treatment creates germ free-like phenotypes in both male and female mice. **a-d**, CFU analysis showed that antibiotic oral gavage depleted gut bacteria. n=3-5 mice. **e-h**, Representative images and bar graph show that antibiotic treatment enlarged the cecum size in male and female mice. Mean ± SEM from 4 mice. **i** and **j**, Bar graph shows the effects of antibiotics and PG muropeptide treatment on spleen size in both male and female mice. n=3-4 mice. *** p<0.001, ** p<0.01, * p<0.05, ns: not significant.

**Extended Data Fig. 7.**
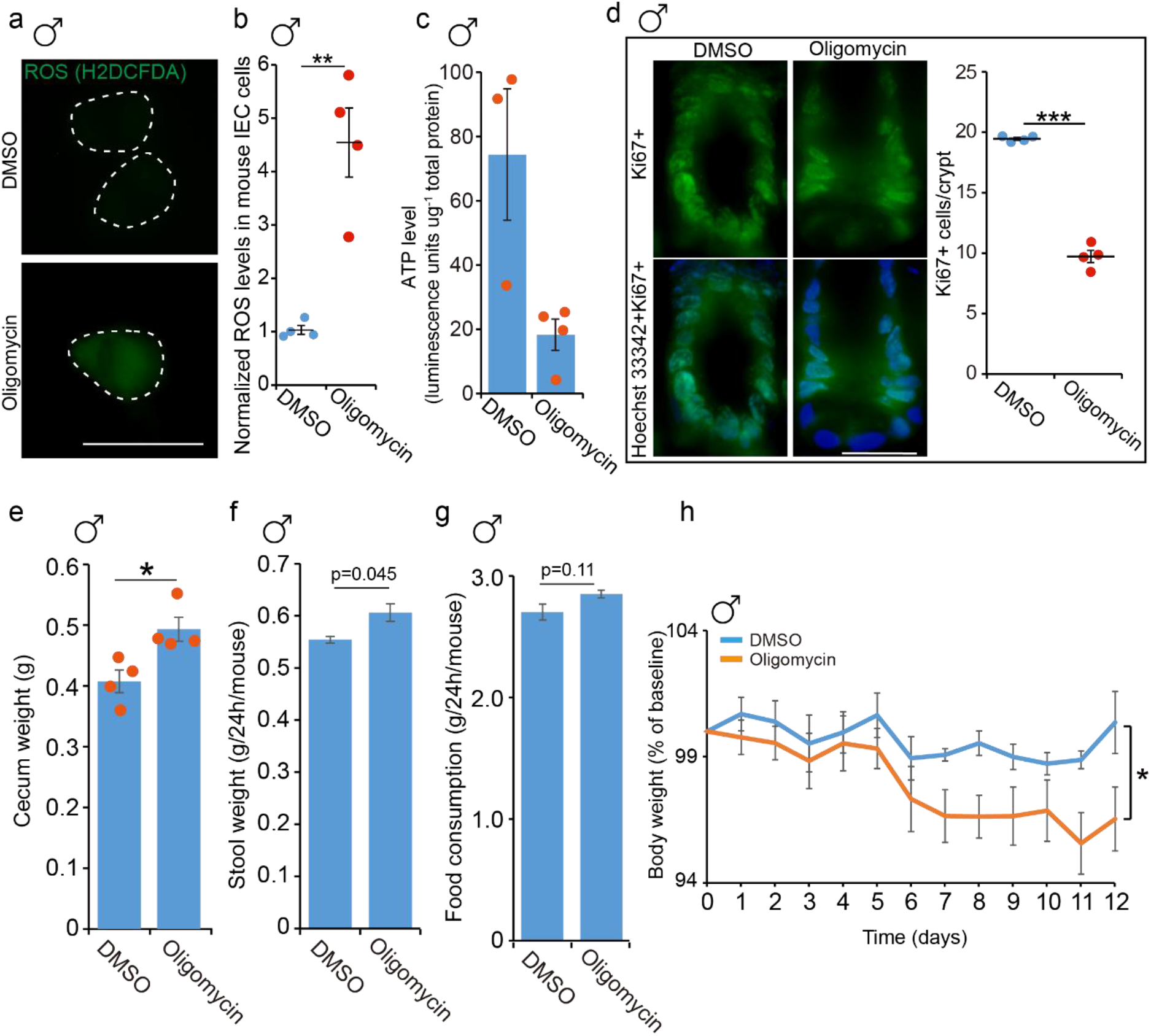
Oligomycin treatment in male mice mimics PG muropeptide depletion induced phenotypes. **a** and **b**, Representative images and quantification showing ROS levels in small IECs of DMSO or oligomycin treated male mice. Mean ± SEM from 4 mice. Scale bar =20 μm. **c**, Bar graph shows ATP level in small IECs of DMSO or oligomycin treated male mice. Mean ± SEM from 3-4 mice. **d**, Representative images and quantification showing proliferating cells (Ki67+) in small intestinal crypts were decreased in Oligomycin-treated male mice, compared to DMSO (mock) treatment. Mean ± SEM from 4 mice. Scale bar =20 μm. **e**, Bar graph showing that oligomycin treatment enlarged the cecum size in male mice. Mean ± SEM from 4 mice. **f**, Quantification showing stool weight was increased in oligomycin treated male mice. Mean ± SEM from 4 mice per 24h. **g**, Quantification showing that oligomycin has no significant effects on food consumption. Mean ± SEM from 4 mice per 24h. **h**, Male mice weight gain during mock (DMSO) or oligomycin oral gavage. ** p<0.01, * p<0.05. ns: not significant.

