## Supplementary material for "Bacterial muropeptides promote OXPHOS and suppress mitochondrial stress in normal and human mitochondrial disease models": Method

**Online contents**

**Method details**

**Cell culture**

The Mode-K cells were a gift from Dr. Karen Madsen at the University of Alberta. The Caco-2 cells were purchased from ATCC (#HTB-37). Cell lines were only used for less than 2 months after thaw. Mode-K cells were maintained in DMEM media (Gibco, 12439047) supplemented with 10% FBS (Gibco, 26140-079), Penicillin (100 U/ml) and Streptomycin (100 μg/ml). Caco-2 cells were maintained in DMEM media (Gibco, 11995040) supplemented with 10% heat-inactivated FBS (Gibco, 26140079), Penicillin (100 U/ml), Streptomycin (100 μg/ml) and 1x non-essential amino acid (Gibco, 11140-050 100x). The fibroblast cells were purchased from Coriell Institute (GM13411) or ATCC (CRL-2091). Fibroblast cells were cultured for less than total 15 passages. Cells were maintained in EMEM media (ATCC, 30-2003) supplemented with 15% FBS (Gibco, 26140-079), Penicillin (100 U/ml) and Streptomycin (100 μg/ml). The cells were maintained in a humidified atmosphere at 37°C, 5% CO_2_.

**Mice**

Experiments were performed after prior approval by the Institutional Animal Care and Use Committee (IACUC) at the University of Colorado Boulder. C57BL/6J strain mice were purchased from Jackson Laboratories and housed in a barrier facility. Animals 6-10 weeks of age, both males and females, were used as noted in figure legends.

**Antibiotic-induced microbiome depletion (AIMD) and PG oral gavage**

For chemical treatment, age-matched mice were randomly assigned to experimental groups, 3-5 mice were housed in each cage. We followed an established protocol to perform antibiotic-induced microbiome depletion (AIMD)^1^. Briefly, mice were given oral gavage of antibiotic cocktail or antibiotic cocktail plus purified peptidoglycan (2 mg/mouse) every 12 h. The antibiotic cocktail contained ampicillin (100 mg/kg), vancomycin (50 mg/kg), metronidazole (100 mg/kg), neomycin (100 mg/kg) and amphotericin B (1mg/kg). The control group received water. The cocktail was freshly made every 24 h. Oral gavage treatment was performed for 23-27 days.

**Oligomycin oral gavage**

Age-matched mice were randomly assigned to experimental groups, 4 mice were housed in each cage. Starting at day 1, mice were given vehicle (0.1% DMSO in PBS) or oligomycin (0.25 mg/kg)^2^ twice a day by oral gavage. Oral gavage treatment was performed for 12-14 days.

**Stool culture measurement**

To assess microbiome depletion by the AIMD method, fresh fecal samples were collected in microcentrifuge tubes containing 1ml sterile PBS. Fecal samples were collected 2 weeks after AIMD initiation. Samples were thoroughly vortexed and diluted with sterile PBS to 1:100. Equal concentrations of stool diluent from different groups were spread on LB agar plates and incubated for 2-3 days at 37 °C. Colony forming units (CFUs) were counted.

**Food consumption and stool weight**

When measuring the food consumption and stool weight, mice in each group were housed in one cage. Food consumption and stool weight were monitored continuously for 3 days. Consumed food and produced stool were weighed every 24 h.

**Cecum and spleen weight**

Mice were euthanized by carbon dioxide asphyxiation followed by cervical dislocation. Cecum and spleen from each mouse were carefully dissected out and weighed by a precision scale.

**Immunohistochemistry**

Small intestines of sacrificed mice were dissected and rapidly flushed with ice cold PBS. A 1.5 cm piece of the middle small intestine (jejunum) was cut off and immediately fixed in 4% paraformaldehyde for 24 h at 4°C before being embedded in optimal cutting temperature (OCT) compound. Sections and slices were cut and prepared by the histology core at the University of Colorado Anschutz campus. After drying at room temperature overnight and heat-induced epitope retrieval in 10 mM sodium citrate (pH 6.0), the sections were blocked 1h at room temperature with IHC/ICC Blocking Buffer (Fisher Scientific, 50-184-85). The sections were incubated 18 h with rabbit anti-Ki67 (Abcam, ab15580) at 4°C. After PBS wash, sections were incubated 2 h with Alexa Fluor 488 conjugated goat anti-rabbit IgG (Fisher Scientific, A11008) at room temperature. After PBS wash, sections were stained with Hoechst dye before being rinsed in water and mounted with mounting medium (Sigma, F4680) under cover slips.

**IEC cell isolation**

A modified protocol^3^ was used to isolate IEC cells from a C57BL/6 mouse. Briefly, animals were euthanized by carbon dioxide asphyxiation followed by cervical dislocation, then the small intestines were dissected. The small intestines (jejunum) were flushed with sterile PBS, opened longitudinally, and cut into 5-mm fragments. The epithelial integrity was disrupted by 1.5 mM dithiothreitol (DTT) treatment on a shaker for 45 min at 37 °C. Liberated IEC cells were collected and separated by Percoll gradient (Sigma, P4937). The interface cells were then collected and used as IEC cells.

**PG isolation, enzyme treatment and supplementation assays**

An adapted method was used to isolate peptidoglycan from bacterial cultures ^4^. In brief, bacterial pellets were resuspended in 1/10 vol 1M NaCl solution and boiled at 100 °C in a heating block for 1h. After washing 4 times with distilled water, the insoluble cell wall preparations were incubated in an ultrasonic water bath (Laboratory Supplies Co., INC, G112SP1T) for 1h, followed by stepwise digestion using DNAse/RNAse and trypsin at 37 °C for 1h, respectively. The solutions were boiled for 5 min at 100°C to inactive the enzymes, followed by washing twice with distilled water. The PG pellets were recovered by centrifugation at 13,000 rpm for 10 min and stored at 4°C.

For further enzymatic treatment of isolated PG, lysozyme (Fisher [44-031-GM](https://www.fishersci.com/shop/products/lysozyme-chicken-egg-white/44031gm#?keyword=true), 300 μg/ml) was added to PG solution (20 mM HEPES pH7.5) and incubated at 37 °C for 48h. The supernatants were collected after centrifugation at 12,000 rpm for 10min and stored at -20°C for later assays.

For Protease-K treatment, 400 μg/ml Protease-K was added to PG solution (20 mM HEPES pH7.5) and incubated at 37°C for 2h. The enzymatic reaction was stopped by boiling at 100°C for 15 min.

**Molecular test**

Total DNA was extracted from wild type fibroblasts or GM13411 cells by DNeasy Blood & Tissue Kits (Qiagen, 69504). To detect the T8993G mutation, fragments of mtDNA were amplified by PCR (Forward primer: CCGACTAATCACCACCCAAC; Reverse primer: TGTCGTGCAGGTAGAGGCTT). The mutation was detected by restriction endonuclease AvaI treatment. Uncut one fragment is wild type 8993T and two cut fragments represent 8993G.

**Recombinant protein expression and purification**

The ATP5H, ATP-2, ATP-4 and ATP-5 cDNA were PCR amplified from HeLa cells or N2 worm cDNA libraries. ATP5A1 and ATPB recombinant protein were purchased from Aviva Systems Biology. All gene constructs were inserted into a pET28a vector. These constructs were transformed into *E. coli* BL21(DE3) cells and grown in LB broth at 37℃ overnight. The overnight cultures were diluted 1:100 vol of fresh LB broth and cultured until OD_600_ reached ~0.6. The expression of recombinant proteins was induced with 0.1 mM IPTG at 20 ℃ for 19 h. Bacterial cells were harvested by centrifugation at 5000xg for 10 min at 4°C. After centrifugation, the pellets were lysed in buffer containing 25 mM HEPES pH 7.5, 500 mM NaCl, 0.5% NP-40, 1 mM DTT and 1x protease inhibitor, followed by centrifugation at 12,000xg for 30 min at 4°C. The supernatants were incubated with Ni-NTA agarose beads (Qiagen, 151028822) for 1 h and rinsed 3 times with a wash buffer (25 mM HEPES pH 7.5, 500 mM NaCl, 1 mM DTT and 30 mM imidazole). Proteins were eluted with buffer containing 25 mM HEPES pH 7.5, 500 mM NaCl, 1mM DTT and 500 mM imidazole. Amicon Ultra centrifugal filters (10kD) were used to concentrate the proteins, followed by elution in buffer (25 mM HEPES pH 7.5, 250 mM NaCl and 1 mM DTT). These proteins were then stored at -80°C with 10% glycerol.

**Verification of the interaction between PG and ATP synthase subunits**

To confirm the interaction between PG and ATP synthase, PG pull-down assays were performed. Total protein from caco-2 cell lysates or recombinant ATP synthase subunits were incubated with PG. Western blots (Invitrogen, 15H4C4) were performed to detect ATP5A1 (caco-2 cells) or His-tag.

**Mitochondria extraction and HEK-Blue NOD1**

Mitochondria from isolated intestinal epithelial cells (IEC) and Mode-K cells were extracted with the Mt Isolation Kit for Cultured Cells (Thermo Scientific, 89874) and Mt from worms were extracted from L4 larvae with the Mt Isolation Kit for Tissue (Thermo Scientific, 89801), following the manufacturer’s protocols.

Isolated Mitochondria were lysed with Tris-buffed saline (25 mM Tris-HCl, pH 7.2, 150 mM NaCl, 1x protease inhibitor) and centrifuged at 12,000xg for 5 min to collect the supernatant. Samples were adjusted based on total Mt protein as determined by the BCA protein assay kit (Thermo Scientific, 23225). HEK293 cells expressing the human NOD1 receptor and the NF-κB SEAP reporter genes (Invitrogen, hkb-hnod1) were used according to the manufacturer’s instructions to assess the muropeptides in mitochondrial lysates. 20 μL of adjusted mitochondria lysates, or lysis buffer (mock), were added to a 96-well plate first. Then, a cell suspension (~280,000 cells per ml in HEK Blue detection medium) was prepared and 180 µl was added to each well with mitochondrial lysates or mock lysate. The 96-well plate was incubated at 37°C, 5% CO_2_ overnight and assessed by reading OD at 650 nm.

**Blue native PAGE analysis of ATP synthase complex**

Mitochondria were extracted from HEK293T cells by kit (ThermoFisher, 89874) or purchased from abcam (ab110338, bovine heart mitochondria) and solubilized by detergent and 1x mitochondrial buffer (abcam, ab109907). After centrifugation, the supernatant was collected for mock or muropeptide treatment. 30 μl mock buffer (lysozyme solution) or muropeptides were mixed with 30 μl mitochondrial supernatant and incubated on ice for 30min. Each sample was then loaded onto a BN-PAGE gel to separate the mitochondrial complexes. After membrane transfer and blocking, the antibody against ATP5A1 (Invitrogen, 15H4C4) was used to detect ATP synthase complex.

**Complex V activity assay**

The complex V activity assay was performed using the MitoTox™ Complex V OXPHOS Activity Assay Kit (Abcam, ab109907), following the manufacturer’s protocol. Approximate 21 μg of mitochondria were added to each well. OD_340_ values were monitored by a SpectraMax M5 plate reader in kinetic mode at 30 °C.

**Measurement of oxygen consumption rates (OCR)**

OCR measurements were performed with a Seahorse XF-24 analyzer (Seahorse Bioscience). Cells were seeded in a Seahorse XF-24 cell culture microplate [1.7 x10^5^ caco-2 cells per well in 250 μl DMEM media (Gibco, 11995040) supplemented with 10% heat-inactivated FBS and 1% non-essential amino acid, or 1.0 x10^4^ fibroblast cells per well in 200 μl EMEM media supplemented with 15% FBS] and then incubated at 37°C and 5% CO_2_ overnight. After overnight recovery, the cells were incubated with muropeptide solution or lysozyme solution (mock) for 24 h. Cells were washed in Seahorse XF DMEM media (Agilent, 30119005, supplemented with 1mM pyruvate, 10 mM glucose and 2 mM glutamine) and incubated at 37°C with no CO_2_ for 1 h before measurement. Cells were washed again with Seahorse XF DMEM media just before measuring. OCR was measured and analyzed by adding 2.5 μM oligomycin, 1μM FCCP and 1.25 μM ROT/AA (Agilent, 103015-100) sequentially following the manufacture’s instruction.

**Measurement of ATP levels**

Cells were seeded in 96-well plates (caco-2 cells: 2x10^4^ cells/well, fibroblasts: 1x10^4^ cells/well). After overnight recovery, cells were treated with either a lysozyme solution (mock) or muropeptides for 24 h. ATP levels were measured using the Luminescent ATP Detection Assay Kit (Abcam, ab113849). Luminescence was monitored by a SpectraMax M5 plate reader. The protein concentrations were measured by a BCA Protein Assay Kit (Thermo Scientific,	23227). ATP levels were normalized to total protein content.

For IEC ATP measurement, IECs were scratched off from small intestine (jejunum) by glass slides. IECs were resuspended in sterile PBS and loaded into a 96-well plate and subjected to ATP Detection and BCA protein assay as described above. ATP levels were normalized to total protein content.

**ROS measurement**

The ROS staining protocol was modified from ^5^. Briefly, isolated mouse IECs or cultured cells were washed three times with MEM (no FBS). Then the cells were transferred into the staining solution (1 uM H2DCFDA in MEM, no FBS) and incubated at 37°C, 5% CO_2_ for 30 min. After washing 4 times with MEM (no FBS), cells were subjected to microscopic examination.

**Measurement of mitochondrial membrane potential**

Caco-2 cells were seeded in 24-well plate [inserted with glass coverslips (Neuvitro, GG12PDL)] at 1x10^4^ cells/well. After overnight recovery, cells were treated with either the lysozyme solution (mock) or muropeptides (800 μg/ml) for 48 h. Culture media was changed every 24 h. MitoTracker-Red CMXRos (Thermo Fisher Scientific, M46752) was diluted into DMEM (Gibco, 11995040) (no FBS) to a final concentration of 100 nM just before use. Caco-2 cells were cultured to 50-60% confluency. The cells were washed 3 times with DMEM (without FBS) and 500 μl MitoTracker-Red CMXRos solution was added. The cell culture plates were then wrapped in foil and maintained at 37 °C and 5% CO_2_ for 30 min. The cells were then washed 3 times with DMEM (no FBS) before microscopic examination. ImageJ was used to analyze mean fluorescence for MitoTracker dyes.

**Transmission electron microscopy**

Cells were grown on sapphire discs and prepared for electron microscopy using high pressure freezing and freeze substitution as described in McDonald et al., 2010. Briefly, 3mm sapphire discs (Technotrade International) were coated with gold and a large F was then scratched into the surface to help orient the cell side. The disks were coated with collagen, sterilized under UV light and cells plated for culturing. Monolayers grown on sapphire discs were frozen using a Wohlwend Compact 02 high pressure freezer (Technotrade International). The frozen cells were then freeze substituted in 1% OsO_4_ and 0.1% uranyl acetate in acetone at -80°C for 3 days then gradually warmed to room temperature. The discs were flat embedded in a thin layer of Epon resin and polymerized at 60°C. Regions containing cells were identified in the light microscope, and a small square of resin containing the cells was excised and remounted onto a blank resin block. Thin (80 nm) sections were cut using a Leica Ultracut UCT and collected onto formvar-coated slot grids. Grids were post stained with 2% uranyl acetate and Reynolds lead citrate. The samples were imaged in a Tecnai T12 Spirit TEM operating at 100kV (Thermo Fisher Scientific, Waltham, MA) using an AMT side-mount CCD camera.

**Western block for Creatine kinase**

For western blot analysis, cultured cells were suspended in lysis buffer containing 25 mM HEPES pH 7.5, 500 mM NaCl, 0.5% NP-40, 1 mM DTT and 1x protease inhibitor, and homogenized by ultrasonication. Samples were diluted with 4x SDS loading buffer and protein were subjected to electrophoresis on 4-12% precast SDS–PAGE gels. Proteins were transferred to PVDF membranes (Millipore, IPVH00010). Thereafter, membranes were incubated in TBST containing 5% milk powder for blocking. Primary antibodies were diluted 1:1,000 and incubated at 4°C overnight. Appropriate HRP-conjugated secondary antibodies were used to detect the respective immunoreactive protein using an enhanced chemiluminescence light-detecting kit (Thermo Scientific, 32109).

**Puromycin incorporation of measurement of protein synthesis**

Puromycin labeling was performed as previously described ^6^. Briefly, cells were incubated in puromycin (MP Biomedicals, 0210055210) containing media (10 μg/ml) for 20 minutes, washed twice with cold PBS, lysed in 1x RIPA buffer (Thermo Scientific, J62725AP) containing proteinase inhibitors and phosphatase inhibitors. 50-100 μg of protein from cell lysate was incubated at 4°C overnight with 1 μg of anti-puromycin antibody (MilliporeSigma, Clone: 12D10). Next, puromycin labeled proteins were purified by MS-Compatible Magnetic IP Kit (Thermo Scientific, 90409) following the manufacturer’s instruction, and processed for mass spectrometry analysis.

**Lactate assay**

Cells were seeded in 24-well culture plate at 4.0 x10^4^ cells/well and incubated at 37°C and 5% CO_2_ overnight. The cells were incubated with muropeptide solution or lysozyme solution (mock) for 24 h before collecting the growth media. Lactate levels in the growth media samples were calculated using the L-Lactate Assay Kit (Cayman Chemical, 700510) according to the manual provided. The lactate levels were normalized by total cell numbers.

**Ammonia/ammonium assay**

Cells were seeded in a 24-well culture plate at 4.0 x10^4^ cells/well and incubated at 37 °C and 5% CO_2_ overnight. The cells were incubated with muropeptide solution or lysozyme solution (mock) for 24 h before collecting the growth media. Ammonia levels in the growth media samples were calculated using the Ammonia/ammonium Assay kit (Bioassay Systems, ENH3-100) according to the manual provided. The ammonia levels were normalized by total cell number.

**Cell viability Assay**

Fibroblast cells were seeded in 24-well plates at 4x10^4^ cells/well. After overnight recovery, cells were treated with either a lysozyme solution (mock) or muropeptides for 24 h. To prepare cell suspension, fibroblast cells were dissociated from culture surfaces by 0.25% Trypsin-EDTA (Gibco, 25200056) and complete growth media was used to inactivate trypsin. A hemocytometer was used to count the cell numbers.

**Microscopy**

Analysis of fluorescence was performed under Nomarski optics on a Zeisis Axioplan2 microscope with a Zeiss AxioCam MRm CCD camera. Plate phenotypes were observed using a Leica MZ16F dissecting microscope with a Hamamatsu C4742-95 CCD camera.

**Quantification and Statistical analysis**

ImageJ software was used to quantify fluorescence intensity. We used the Student’s t-test to determine the significant differences in all figures.

1	Zarrinpar, A. *et al.* Antibiotic-induced microbiome depletion alters metabolic homeostasis by affecting gut signaling and colonic metabolism. *Nature communications* **9**, 2872, doi:10.1038/s41467-018-05336-9 (2018).

2	Franchi, L. *et al.* Inhibiting Oxidative Phosphorylation In Vivo Restrains Th17 Effector Responses and Ameliorates Murine Colitis. *Journal of immunology* **198**, 2735-2746, doi:10.4049/jimmunol.1600810 (2017).

3	Roulis, M., Armaka, M., Manoloukos, M., Apostolaki, M. & Kollias, G. Intestinal epithelial cells as producers but not targets of chronic TNF suffice to cause murine Crohn-like pathology. *Proceedings of the National Academy of Sciences of the United States of America* **108**, 5396-5401, doi:10.1073/pnas.1007811108 (2011).

4	Kuhner, D., Stahl, M., Demircioglu, D. D. & Bertsche, U. From cells to muropeptide structures in 24 h: peptidoglycan mapping by UPLC-MS. *Scientific reports* **4**, 7494, doi:10.1038/srep07494 (2014).

5	Wu, D. & Yotnda, P. Production and detection of reactive oxygen species (ROS) in cancers. *Journal of visualized experiments : JoVE*, doi:10.3791/3357 (2011).

6	Schmidt, E. K., Clavarino, G., Ceppi, M. & Pierre, P. SUnSET, a nonradioactive method to monitor protein synthesis. *Nature methods* **6**, 275-277, doi:10.1038/nmeth.1314 (2009).
